# Wild Patagonian yeast improve the evolutionary potential of novel interspecific hybrid strains for Lager brewing

**DOI:** 10.1101/2024.01.29.577692

**Authors:** Jennifer Molinet, Juan P. Navarrete, Carlos A. Villarroel, Pablo Villarreal, Felipe I. Sandoval, Roberto F. Nespolo, Rike Stelkens, Francisco A. Cubillos

**Author notes:** Address correspondence to Francisco A. Cubillos, and Jennifer Molinet.

## Abstract

Lager yeasts are limited to a few strains worldwide, imposing restrictions on flavour and aroma diversity and hindering our understanding of the complex evolutionary mechanisms during yeast domestication. The recent finding of diverse *S. eubayanus* lineages from Patagonia offers potential for generating new Lager yeasts and obtaining insights into the domestication process. Here, we leverage the natural genetic diversity of *S. eubayanus* and expand the Lager yeast repertoire by including three distinct Patagonian *S. eubayanus* lineages. We used experimental evolution and selection on desirable traits to enhance the fermentation profiles of novel *S. cerevisiae* x *S. eubayanus* hybrids. Our analyses reveal an intricate interplay of pre-existing diversity, selection on species-specific mitochondria, *de-novo* mutations, and gene copy variations in sugar metabolism genes, resulting in high ethanol production and unique aroma profiles. Hybrids with *S. eubayanus* mitochondria exhibited greater evolutionary potential and superior fitness post-evolution, analogous to commercial Lager hybrids. Using genome-wide screens of the parental subgenomes, we identified genetic changes in *IRA2*, *SNF3*, *IMA1* and *MALX* genes that influence maltose metabolism, and increase glycolytic flux and sugar consumption in the evolved hybrids. Functional validation and transcriptome analyses confirmed increased maltose-related gene expression, influencing grater maltotriose consumption in evolved hybrids. This study demonstrates the potential for generating industrially viable Lager yeast hybrids from wild Patagonian strains. Our hybridization, evolution, and mitochondrial selection approach produced hybrids with high fermentation capacity, expands Lager beer brewing options, and deepens our knowledge of Lager yeast domestication.

## INTRODUCTION

Humans have paved the way for microbes, such as yeast, to evolve desirable features for making bread, wine, beer and many other fermented beverages for millennia (Steensels et al., 2019). The fermentation environment, characterized by limited oxygen, high ethanol concentrations, and microbial competition for nutrients (typically yeasts, molds, and bacteria) can be considered stressful (Walker & Basso, 2019). One evolutionary mechanism to overcome harsh conditions is hybridization, because it rapidly combines beneficial phenotypic features of distantly related species and generates large amounts of genetic variation available for natural selection to act on (Abbott et al., 2013; Steensels et al., 2021; R. Stelkens & Bendixsen, 2022). Hybrids can also express unique phenotypic traits not seen in the parental populations through the recombination of parental genetic material, enabling them to thrive in different ecological niches (Rieseberg et al., 2003; Steensels et al., 2021; R. B. Stelkens et al., 2014; R. Stelkens & Seehausen, 2009). An iconic example is the domestication of the hybrid yeast *Saccharomyces pastorianus* to produce modern lager (pilsner) beers. *S. pastorianus* is the result of the successful interspecies hybridization between *S. cerevisiae* and *S. eubayanus* (Gallone et al., 2019; Langdon et al., 2019). Hybrids have been shown to benefit from the cold tolerance of *S. eubayanus* and the superior fermentation kinetics of *S. cerevisiae* (Gibson et al., 2017). We now know that domestication over the last 500 years has generated Lager yeast strains with the unique ability to rapidly ferment at lower temperatures resulting in a crisp flavour profile and efficient sedimentation, improving the clarity of the final product. However, the genetic diversity of commercial Lager yeast strains is extremely limited, mainly due to the standardization of industrial Lager production during the nineteenth century in Germany (Gallone et al., 2019; Hutzler et al., 2023). This gave rise to only two genetically distinct *S. pastorianus* subgroups, Group 1 strains (‘Saaz’) and Group 2 strains (‘Frohberg’). The poor genetic diversity of Lager strains used in commercial brewing today (85 Lager strains commercially available versus 358 Ale strains (Bonatto, 2021)) puts tight constraints on the variety of flavours and aromas found in Lager beer. At the same time, it limits our understanding of the evolutionary mechanism operating during the yeast domestication process.

The discovery of *S. eubayanus* in Patagonia in 2011 (Libkind et al., 2011), opened new possibilities for creating novel hybrid strains by using the full range of natural genetic diversity found in this species. It also provides an opportunity to better understand the lager yeast domestication process. Phylogenetic analyses have revealed six distinct lineages of *S. eubayanus*, including China, Patagonia A (‘PA’), Holarctic, and Patagonia B, ‘PB-1’, ‘PB-2’ and ‘PB-3’, and some admixed strains derived from ancient crosses (Langdon et al., 2020; Nespolo et al., 2020). Of these, *S. eubayanus* from Patagonia displays the broadest phenotypic diversity for a wide range of traits, including high maltose consumption, aroma profiles and fermentation capacity (Mardones et al., 2020; Nespolo et al., 2020; Urbina et al., 2020). The distinctive traits of wild Patagonian *S. eubayanus* strains indicate their potential for crafting new Lager beer styles. These strains could yield novel taste and aroma profiles, approaching similar complexity and diversity in flavour, appearance, and mouthfeel as Ale beers.

Lager yeast hybrids experienced an intense domestication process through selection and re-pitching during beer fermentation since the 17^th^ century (Gallone et al., 2019; Gorter De Vries, Pronk, et al., 2019; Hutzler et al., 2023; Langdon et al., 2019; Okuno et al., 2015), a process similar to experimental evolution (Gibson et al., 2020; Gorter De Vries, Voskamp, et al., 2019). Experimental evolution with microbes is a powerful tool to study adaptive responses to selection under environmental constraints (Barrick & Lenski, 2013; Cooper, 2018; Maddamsetti et al., 2015; Payen & Dunham, 2016). Recent studies on novel *S. cerevisiae* x *S. eubayanus* hybrids suggest that hybrid fermentative vigour at low temperature results from a variety of genetic changes, including loss of heterozygosity (LOH), ectopic recombination, transcriptional rewiring, selection of superior parental alleles (Sipiczki, 2018), heterozygote advantage due to the complementation of loss-of-function mutations in the F1 hybrid genome (Brouwers et al., 2019), and novel structural and single nucleotide variants in the hybrid genome (Krogerus, Holmström, et al., 2018). A recent transcriptome analyses of a laboratory-made Lager hybrid strain under fermentation conditions highlighted that the regulatory ‘cross-talk’ between the parental subgenomes caused a novel sugar consumption phenotype in the hybrid (maltotriose utilization, essential for Lager fermentation), which was absent in both parental strains (Brouwers et al., 2019). Although these studies have greatly contributed to our understanding of the genetic basis of different lager phenotypes, most studies only considered a single *S. eubayanus* genetic background (type strain CBS 12357), which alone is not representative of the species-rich genetic diversity.

Here, we hybridized three different *S. cerevisiae* and *S. eubayanus* strains to generate genetically and phenotypically diverse novel Lager hybrids via spore-to-spore mating. The initial *de novo* hybrids had fermentation capacities comparable to those of their parental strains and did not show positive heterosis. However, when we subjected hybrids to a ‘fast motion’ improvement process using experimental evolution under different fermentation conditions for 250 generations, they exceeded the fitness of the ancestral hybrids, particularly those retaining the *S. eubayanus* mitochondria. Superior hybrid fitness was explained by faster fermentation performance and greater maltose and maltotriose consumption We found that copy number variation in *MAL* genes in the *S. cerevisiae* subgenome, together with SNPs in genes related to glycolytic flux, induced significantly greater expression levels of *MAL* and *IMA1* genes, and led to improved fitness under fermentative conditions in these novel *S. cerevisiae* x *S. eubayanus* yeast hybrids. Furthermore, evolved hybrids had significantly distinct aroma profiles, varying significantly from the established profiles found in lager beer.

## MATERIALS AND METHODS

### Parental strains

Three *S. cerevisiae* strains were selected for hybridization from a collection of 15 strains isolated from different wine-producing areas in Central Chile and previously described by (Martinez et al., 2004). Similarly, three *S. eubayanus* parental strains were selected from a collection of strains isolated from different locations in Chilean Patagonia, exhibiting high fermentative capacity and representative of the different Patagonia-B lineages (PB-1, PB-2 and PB-3) (Nespolo et al., 2020). The *S. pastorianus* Saflager W34/70 (Fermentis, France) strain was used as a commercial Lager fermentation control. All strains were maintained in YPD agar (1% yeast extract, 2% peptone, 2% glucose and 2 % agar) and stored at -80 °C in 20% glycerol stocks. Strains are listed in **Table S1A**.

### Interspecific hybrids strains and mitochondria genotyping

Parental strains were sporulated on 2% potassium acetate agar plates (2% agar) for at least seven days at 20 °C. Interspecific F1 hybrids were generated through spore-spore mating between *S. eubayanus* strains and *S. cerevisiae* strains (**Figure S1**). For this, tetrads were treated with 10 μL Zymolyase 100 T (50 mg/mL) and spores of opposite species were dissected and placed next to each other on a YPD agar plates using a SporePlay micromanipulator (Singer Instruments, UK). Plates were incubated at two different temperatures, 12 and 20 °C, for 2-5 days to preserve the cold- and heat-tolerant mitochondria, respectively, as previously described (Baker et al., 2019; Hewitt et al., 2020), resulting in nine different F1 hybrids (ranging from H1 until H9, **Table S1A**). This procedure was repeated on 25 tetrads of each species, for each type of cross (H1 to H9) and temperature (12 and 20 °C), resulting in 18 different cross x temperature combinations. Finally, colonies were isolated, re-streaked on fresh YPD agar plates, and continued to be incubated at 12 and 20 °C. The hybrid status of isolated colonies was confirmed by amplification of rDNA-PCR (ITS1, 5.8S and ITS2) using universal fungal primers ITS1 and ITS4 (Esteve-Zarzoso et al., 1999), followed by digestion of the amplicon using the *Hae*III restriction enzyme (Promega, USA) as previously described (Krogerus et al., 2016) on one colony for each cross attempt (**Figure S1**). Confirmed F1 hybrids were designated as H1 to H9 based on parental strains, followed by the hybridization temperature (12 or 20) and the colony number (i.e. H1.20-1 depicts cross 1 at 20 °C (**Table S1A**)). We identified the mitochondrial genotype by Sanger sequencing the mitochondrial *COX3* gene as previously described (Hewitt et al., 2020).

### Beer wort fermentation and metabolite screening

Fermentations were carried out in three biological replicates using previously oxygenated (15 mg/L) 12 °P wort, supplemented with 0.3 ppm ZnCl_2_ as previously described (Mardones et al., 2020). Briefly, pre-cultures were grown in 5 mL 6 °P wort for 24 h at 20 °C with constant agitation at 150 rpm. Cells were then transferred to 50 mL 12 °P wort and incubated for 24 h at 20 °C with constant agitation at 150 rpm. Cells were collected by centrifugation and used to calculate the final cell concentration to inoculate the subsequent fermentation according to the formula described by (White & Zainasheff, 2010). Cells were inoculated into 50 mL 12 °P wort in 250 mL bottles covered by airlocks containing 30% glycerol. The fermentations were incubated at 12 or 20 °C, with no agitation for 15 days and monitored by weighing the bottles daily to determine weight loss over time.

Sugar (glucose, fructose, maltose and maltotriose) consumption and ethanol production were determined by High-Performance Liquid Chromatography (HPLC) after 14 days of fermentation. Filtered samples (20 μL) were injected in a Shimadzu Prominence HPLC (Shimadzu, USA) with a BioRad HPX-87H column using 5 mM sulfuric acid and 4 mL acetonitrile per liter of sulfuric acid as the mobile phase at a 0.5 mL/min flow rate. Volatile compound production was determined by using HeadSpace Solid-Phase MicroExtraction followed by Gas Chromatography-Mass Spectrometry (HS-SPME-GC/MS) after 14 days of fermentation as previously described (Urbina et al., 2020).

### Phenotypic characterization

Hybrids and parental strains were phenotypically characterized under microculture conditions as previously described (Molinet, Urbina, et al., 2022). Briefly, we estimated mitotic growth in 96-well plates containing Yeast Nitrogen Base (YNB) supplemented with 2% glucose, 2% maltose, 2% maltotriose, 2% glucose and 9% ethanol, 2% glucose and 10% sorbitol, and under carbon source switching from glucose to maltose as previously described (Molinet, Eizaguirre, et al., 2022). All conditions were evaluated at 25 °C. Lag phase, growth efficiency and the maximum specific growth rate were determined as previously described (Ibstedt et al., 2015; Warringer & Blomberg, 2003). For this, the parameters were calculated following curve fitting (OD values were transformed to ln) using the Gompertz function (Zwietering M et al., 1990) in R (version 4.03).

Mid-parent and best-parent heterosis were determined as previously described (Bernardes et al., 2017; Steensels et al., 2014), using equation 1 and 2, where mid-parent heterosis denotes the hybrid deviation from the mid-parent performance and best-parent heterosis denotes the hybrid deviation from the better parent phenotypic value (Zörgö et al., 2012).

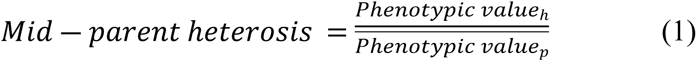

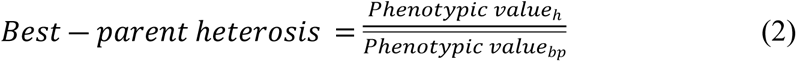

Where:

*Phenotypic value_h_ = Phenotypic value_hybrid_*

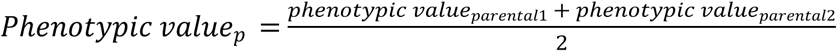

*Phenotypic value_bp_ = max(Phenotypic value_parental_*_1_,*Phenotypic value_parental_*_2_)

### Experimental evolution

Experimental evolution was carried out at 20°C under two different media conditions (M and T): 1) YNB + 2% maltose supplemented with 9% ethanol (M) and 2) YNB + 1% maltose + 1% maltotriose supplemented with 9% ethanol (T). Experimental evolution assays in maltose were performed in a final volume of 1 mL in 2 mL tubes, while those in maltose and maltotriose were performed in a 96-well plate under a final volume of 200 μL. Each hybrid strain was first grown in 0.67% YNB medium with 2% maltose at 25 °C for 24 h with constant agitation at 150 rpm. Each pre-inoculum was then used to inoculate each evolution line at an initial OD_600nm_ of 0.1, with three replicate lines per strain in medium M and four replicate lines in medium T. Lines in medium M were incubated at 20 °C for 72 h. Lines in medium T were incubated for 144 h at 20 °C. After this, cultures were then serially transferred into fresh medium for an initial OD_600nm_ of 0.1. Serial transfers were repeated for 250 generations in total (approximately seven months). The number of generations was determined using the formula log(final cells – initial cells)/log_2_ (Mardones et al., 2021). Population samples were stored at -80 °C in 20% glycerol stocks after 50, 100, 150, 200 and 250 generations. After 250 generations, three colonies were isolated for each replicate line on YPM solid medium (1% yeast extract, 2% peptone, 2% maltose and 2 % agar) supplemented with 6% ethanol. The fastest growing colonies were stored at -80 °C in 20% glycerol stocks. The fitness increase of each the 28 evolved line was determined as the ratio between the phenotypic value of a given line and the equivalent of its respective ancestral hybrid.

### Genomic characterization

Genomic DNA was obtained for whole-genome sequencing using the YeaStar Genomic DNA Kit (Zymo Research, USA) and sequenced in an Illumina NextSeq500 following the manufacturer’s instructions. Variant calling and filtering were done with GATK version 4.3.0.0 (Depristo et al., 2011). Briefly, cleaned reads were mapped to a concatenated reference genome consisting of *S. cerevisiae* strain DBVPG6765 (Yue et al., 2017) and S*. eubayanus* strain CL216.1 (Mardones et al., 2020) using BWA mem 0.7.17 (Li & Durbin, 2010), after which output bam files were sorted and indexed using Samtools 1.13 (Li et al., 2009). Variants were called per sample using HaplotypeCaller (default settings) generating g.vcf files. Variant databases were built using GenomicsDBImport and genotypes were called using GenotypeGVCFs (-G StandardAnnotation). SNPs and INDELs were extracted and filtered out separately using SelectVariants. We then applied recommended filters with the following options: QD < 2.0, FS > 60.0, MQ < 40.0, SOR > 4.0, MQRankSum < -12.5, ReadPosRankSum < -8.0. This vcf file was further filtered removing missing data using the option –max-missing 1, filtering out sites with a coverage below 5^th^ or above the 95^th^ coverage sample percentile using the options –min-meanDP and –max-meanDP, and minimum site quality of 30 (--minQ 30) in vcftools 0.1.16 (Danecek et al., 2011). Sites with a mappability less than 1 calculated by GenMap 1.3.0 (Pockrandt et al., 2020) were filtered using bedtools 2.18 (Quinlan & Hall, 2010). As an additional filtering step, the ancestral and evolved vcf files were intersected using BCFtools 1.3.1 (Danecek et al., 2021) and variants with shared positions were extracted from the vcf files of the evolved hybrids. Annotation and effect prediction of the variants were performed with SnpEff (Cingolani et al., 2012).

We used sppIDer (Langdon et al., 2018) to assess the proportional genomic contribution of each species to the nuclear and mitochondrial genomes in each sequenced hybrid. In addition, we used the tool to identify potential aneuploidies within these genomes. CNVs were called using CNVkit (--method wgs, - -target-avg-size 1000) (Talevich et al., 2016). As the analysis was performed on a haploid reference (both parental genomes were present), a CNV of log2 = 1 corresponds to a duplication.

### RNA-seq analysis

Gene expression analysis was performed on ancestral and evolved hybrid strains H3-A and H3-E. RNA was obtained and processed after 24 h under beer wort fermentation in triplicates, using the E.Z.N.A Total RNA kit I (OMEGA) as previously described (Molinet, Eizaguirre, et al., 2022; Venegas et al., 2023). Total RNA was recovered using the RNA Clean and Concentrator Kit (Zymo Research). RNA integrity was confirmed using a Fragment Analyzer (Agilent). Illumina sequencing was performed in NextSeq500 platform.

Reads quality was evaluated using the fastqc tool (https://www.bioinformatics.babraham.ac.uk/projects/fastqc/) and processed using fastp (-3 l 40) (Chen et al., 2018). Reads were mapped to a concatenated fasta file of the DBVPG6765 and CL216.1 genome sequences. To account for mapping bias due to the different genetic distances of the parental strains to their reference strains, the L3 and CL710.1 parental strains were re-sequenced using WGS, after which genomic reads were mapped with BWA (Li & Durbin, 2010) to the DBVPG6765 and CL216 references and SNPs were called using freebayes (Garrison & Marth, 2012). These SNPs were used to correct the hybrid genome sequence using the GATK FastaAlternateReferenceMaker tool. RNAseq reads were mapped to this hybrid reference using STAR (-outSAMmultNmax 1,-outMultimapperOrder random) (Dobin et al., 2013). Counts were obtained with featureCounts using a concatenated annotation file (Liao et al., 2014). Counts were further analyzed in R using de DESeq package (Love et al., 2014). A PCA analysis to evaluate the reproducibility of replicates was performed, after which two outlier replicates (H3-A replicate 3 and H3-E replicate 2) were removed. To analyze differences in allele expression, a list of 1-to-1 orthologous genes between both parental strains were identified using OMA (Zahn-Zabal et al., 2020). Orthologous genes that differ more than 5% on their gene lengths were excluded. The differential allelic expression of these orthologous genes was determined using design = ∼parental, with parental being “L3” or “CL710”. Furthermore, orthologous genes that showed differential allele expression depending on the ancestral or evolved strain background were assessed using an interaction term (∼ parental:condition), with condition being “ancestral” or “evolved”. Finally, to evaluate differences between ancestral and evolved hybrid strains, all 11,047 hybrid genes (5,508 *S. eubayanus* and 5,539 *S. cerevisiae*) were individually tested for differential expression using DESeq2. Overall gene expression differences were evaluated using the design ∼condition. For all analyzes an FDR < 0.05 was used to consider statistical differences. GO term enrichment analyzes on differentially expressed genes were calculated using the package TOPGO (Alexa & Rahnenfuhrer, 2023).

### *IRA2* gene validation

The *S. cerevisiae IRA2* polymorphism was validated by Sanger sequencing. PCR products were purified and sequenced by KIGene, Karolinska Institutet (Sweden). The presence of the SNP in the evolved hybrid strains was checked by visual inspection of the electropherograms. Null mutants for the *IRA2* gene in the *S. cerevisiae* sub-genome were generated using CRISPR-Cas9 (Dicarlo et al., 2013) as previously described (Molinet, Urbina, et al., 2022). Briefly, the gRNAs were designed using the Benchling online tool (https://www.benchling.com/) and cloned into the pAEF5 plasmid (Fleiss et al., 2019), using standard “Golden Gate Assembly” (Horwitz et al., 2015). Ancestral and evolved hybrids were co-transformed with the pAEF5 plasmid carrying the gRNA and the Cas9 gene, together with a double-stranded DNA fragment (donor DNA). The donor DNA contained nourseothricin (NAT) resistance cassette, obtained from the pAG25 plasmid (Addgene plasmid #35121), flanked with sequences of the target allele, corresponding to 50-pb upstream of start codon and 50-pb downstream of the stop codon. Correct gene deletion was confirmed by standard colony PCR. All primers, gRNAs, and donor DNA are listed in **Table S1B**.

### Statistical analysis

Data visualization and statistical analyses were performed with R software version 4.03. Maximum specific growth rates and total CO_2_ loss were compared using an analysis of variance (ANOVA) and differences between the mean values of three replicates were tested using Student’s t-test and corrected for multiple comparisons using the Benjamini-Hochberg method. A *p-value* less than 0.05 (*p*<0.05) was considered statistically significant. Heatmaps were generated using the ComplexHeatmap package version 2.6.2. A principal component analysis (PCA) was performed on phenotypic data using the FactoMineR package version 2.4 and the factoextra package version 1.07 for extracting, visualizing and interpreting the results.

### Data availability

All fastq sequences were deposited in the National Center for Biotechnology Information (NCBI) as a Sequence Read Archive under the BioProject accession number PRJNA1043100 (http://www.ncbi.nlm.nih.gov/bioproject/1043100).

## RESULTS

*De novo S. cerevisiae* x *S. eubayanus* F1 hybrids show similar phenotypes as their parental strains.

The *S. cerevisiae* and *S. eubayanus* parental strains were selected from a previously described collection of Chilean isolates by (Martinez et al., 2004) and (Nespolo et al., 2020), respectively (**Table S1A**). Initially, three *S. cerevisiae* strains from vineyards were selected because they showed: i) the highest maximum CO_2_ loss in beer wort (**Figure S2A, Table S2**), ii) the best growth performance under maltotriose conditions (**Figure S2B**), and iii) the most efficient maltotriose uptake during microculture conditions (**Figure S2C)**. These strains were L3, L270, and L348. The selection of *S. eubayanus* parental strains was determined by two criteria: i) to represent distinct lineages found in the Chilean Patagonia to maximize genetic diversity (one strain per lineage, PB-1, PB-2, and PB-3), and ii) to display the highest CO_2_ loss during fermentation when compared to strains within their respective lineages based on previous assays (Nespolo et al., 2020). In this way, we selected CL450.1, CL710.1 and CL216.1, from PB-1, PB-2, and PB-3, respectively.

We first assessed sporulation efficiency and spore viability in the six chosen parental strains (**Table S2C**). Sporulation efficiency ranged from 12.7% to 95.5% and spore viability ranged from 15% to 100% across strains. Then, nine different interspecific F1 hybrid crosses were created by mating three *S. cerevisiae* and three *S. eubayanus* strains through spore-to-spore mating (**Figure S1**). Mating was conducted at 12 °C and 20 °C to promote the preservation of the cold- and heat-tolerant mitochondria, respectively, as previously described (Baker et al., 2019; Hewitt et al., 2020). We obtained 31 interspecific hybrids (**Table S1A**), which we phenotyped individually under microculture conditions. In this way, we estimated microbial growth under similar conditions to those encountered during beer wort fermentation, such as glucose, maltose, maltotriose and ethanol (**Table S3**). Hierarchical clustering of the phenotypic data denotes three main clusters, where there was no discernible clustering of hybrids based on their parental strains or hybridization temperature, highlighting the considerable phenotypic diversity resulting from hybridization (**Figure 1A**). To describe the phenotypic landscape of the 31 hybrids more comprehensively, we conducted a PCA analysis (**Figure 1B**). The individual factor map shows that hybrids made at 20°C fall into the right upper quarter of the phenotype space, and are associated with higher growth rate in media with maltose and glucose compared to hybrids made at 12 °C. This was particularly the case for four hybrid strains (H1, H3, H4 and H6), involving parental strains L3, L270, CL216.1 and CL710.1 (all p-values < 0.05, one-way ANOVA, **Table S3B**).

**Figure 1.**
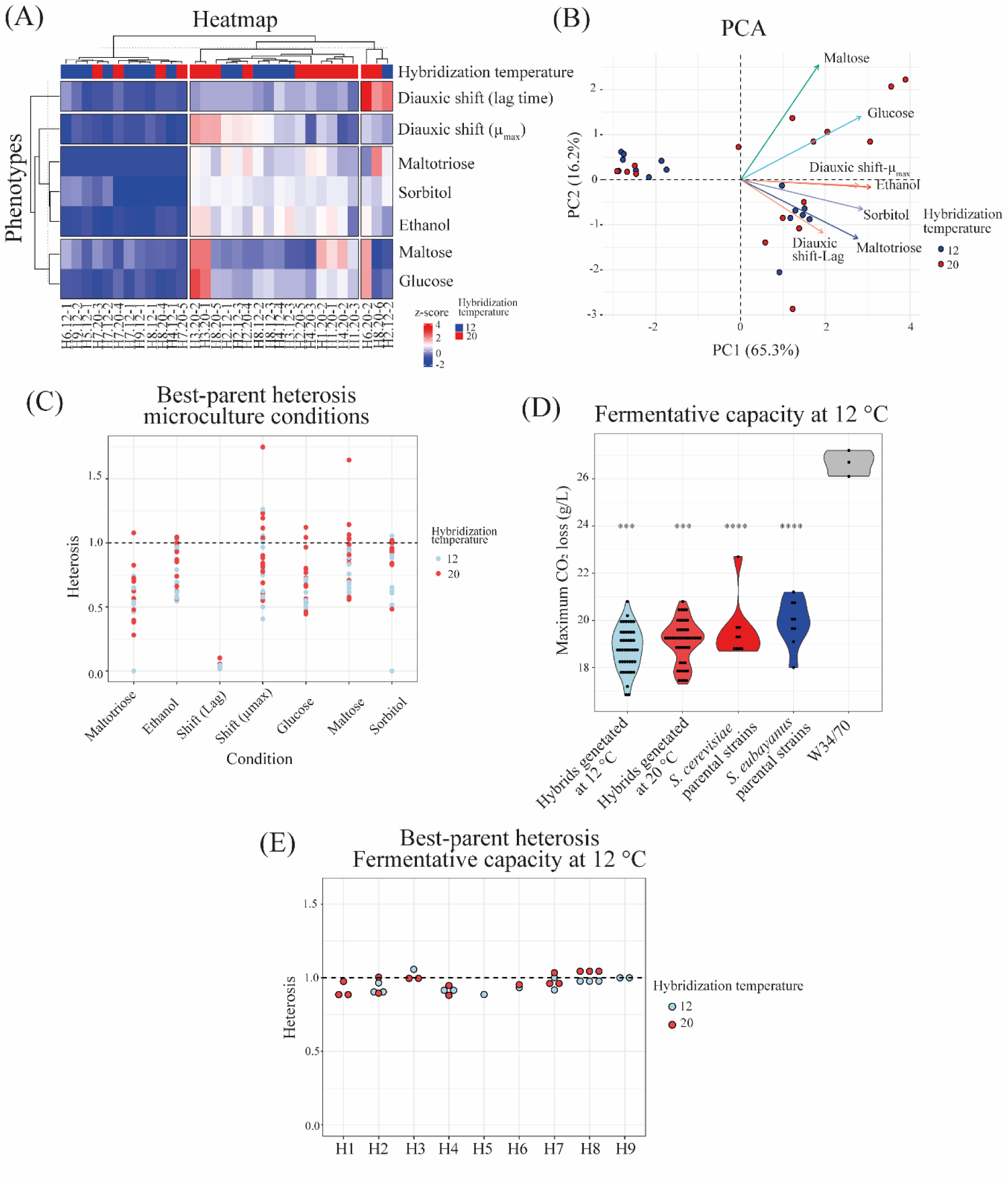
Phenotypic characterization of interspecific F1 hybrids. A) Hierarchically clustered heatmap of phenotypic diversity of 31 interspecific hybrids strains under microculture conditions. Phenotypic values are calculated as normalized z-scores. (B) Principal component analysis (PCA) using the maximum specific growth rates under six microculture growth conditions, together with the distribution of hybrid strains. Arrows depict the different environmental conditions. (C) Best-parent heterosis in the 31 interspecific hybrids evaluated under microculture conditions in triplicates. (D) Fermentation capacity for the 31 interspecific hybrids and parental strains at 12 °C. Plotted values correspond to mean values of three independent replicates for each hybrid. Asterisk indicates different levels of significance compared to the commercial strain W34/70 (Student t-test; *** p≤ 0.001 and **** p≤ 0.0001). (E) Best-parent heterosis in the 31 interspecific hybrids evaluated under fermentation conditions at 12 °C.

To assess the impact of hybridization on yeast fitness, we calculated best-parent and mid-parent heterosis coefficients across the 31 hybrids (**Figure 1C, Table S3C-3D**). While some hybrids exhibited positive mid-parent heterosis in 5 out 7 conditions (**Table S3C**), we generally did not observe hybrids with positive best-parent heterosis (BPH, **Table S3D)**, except for rare cases involving maltose utilization and growth rate during diauxic shift, where 2 and 5 hybrids, respectively, displayed positive values (**Figure 1C**). For example, in the H3.20-1 hybrid (L3 x CL710.1, generated at 20°C) we obtained a 74.8% BPH value for growth rate during diauxic shift. Overall, inter-species hybridization did not result in a significant enhancement of fitness in F1 hybrids.

Considering the potential use of these new hybrids for the production of Lager beer, we proceeded to assess the fermentation capacity of the 31 hybrids in wort at low temperature (**Figure 1D, Table S4**). Hybrids generated at 12 °C displayed similar levels of CO_2_ production compared to those obtained at 20 °C (**Figure 1D, Table S4A,** p-value = 0.17, one-way ANOVA). We did not observe any hybrids exhibiting superior fermentative capacity when compared to their respective parental strains (**Figure S3**), and there was no evidence for hybrid vigour according to best-parent and mid-parent heterosis coefficients (**Figure 1E, Table S4C-D**). Neither parents nor hybrids reached the fermentative capacity of the commercial strain W34/70 (p-value < 0.05, one-way ANOVA).

### Evolved lines carrying the *S. eubayanus* mitochondria exhibit a greater fitness under fermentation

All results so far indicated that the *de novo* interspecific hybrids did not show any hybrid vigour, in none of the phenotypes assessed. We thus decided to subject hybrids to experimental evolution to enhance their fermentative capacity. We specifically selected four hybrids (H3.12-3, H4.12-4, H6.20-2, and H8.20-5) because they demonstrated the highest phenotypic values across kinetic parameters. From here on we will refer to these strains as H3-A, H4-A, H6-A and H8-A (A for ‘ancestral’ or unevolved hybrid). These four hybrids completely consumed the sugars present in the beer wort, except for maltotriose, which may explain the lower fermentative capacity of the hybrids compared to the commercial strain W34/70 (**Table S4E**). Furthermore, these four hybrids represent crosses made at 12 °C and 20 °C and they encompass all six parental genetic backgrounds. To enhance the fermentative capacity of these selected hybrids, they were subjected to adaptive evolution at 20 °C for 250 generations under two distinct conditions: i) YNB supplemented with 2% maltose and 9% ethanol (referred to as “M” medium), and ii) YNB supplemented with 1% maltose, 1% maltotriose, and 9% ethanol (referred to as “T” medium). We evolved three lines independently per cross in medium M, and four independent lines per cross in medium T. These conditions were chosen because maltose is the main sugar in beer wort (approximately 60%) (Nikulin et al., 2018). Considering that yeast typically consume carbon sources in a specific order (glucose, fructose, maltose, and maltotriose), we employed a combination of maltose and maltotriose to facilitate the utilization of the latter carbon source.

After 250 generations, the evolved lines showed different levels of fitness improvements, depending on the environmental conditions and their genetic background (**Figure 2A, Figure S4**), with distinct fitness trajectories over time (**Figure S5**). All evolved lines significantly increased in fitness in at least one of the evolution media and/or kinetic parameters assessed compared to their respective ancestral hybrids (**Figure 2A, Table S5A-B**; p-value < 0.05, one-way ANOVA). Interestingly, evolved lines from hybrids made at 12 °C mating temperature (H3-A and H4-A) showed a more pronounced fitness increase in the T medium compared to those generated at 20 °C (p-value = 3.327e-08, one-way ANOVA, **Figure 2B** and **S4B**), suggesting that hybrids with *S. eubayanus* mitochondria have greater potential for improvement than hybrids with *S. cerevisiae* mitochondria. We verified that the two ancestral H3-A and H4-A hybrids carried only *S. eubayanus* mitochondria by sequencing the *COX3* gene, while H6-A and H8-A inherited the mitochondria from *S. cerevisiae* (**Table S5C-D**).

**Figure 2.**
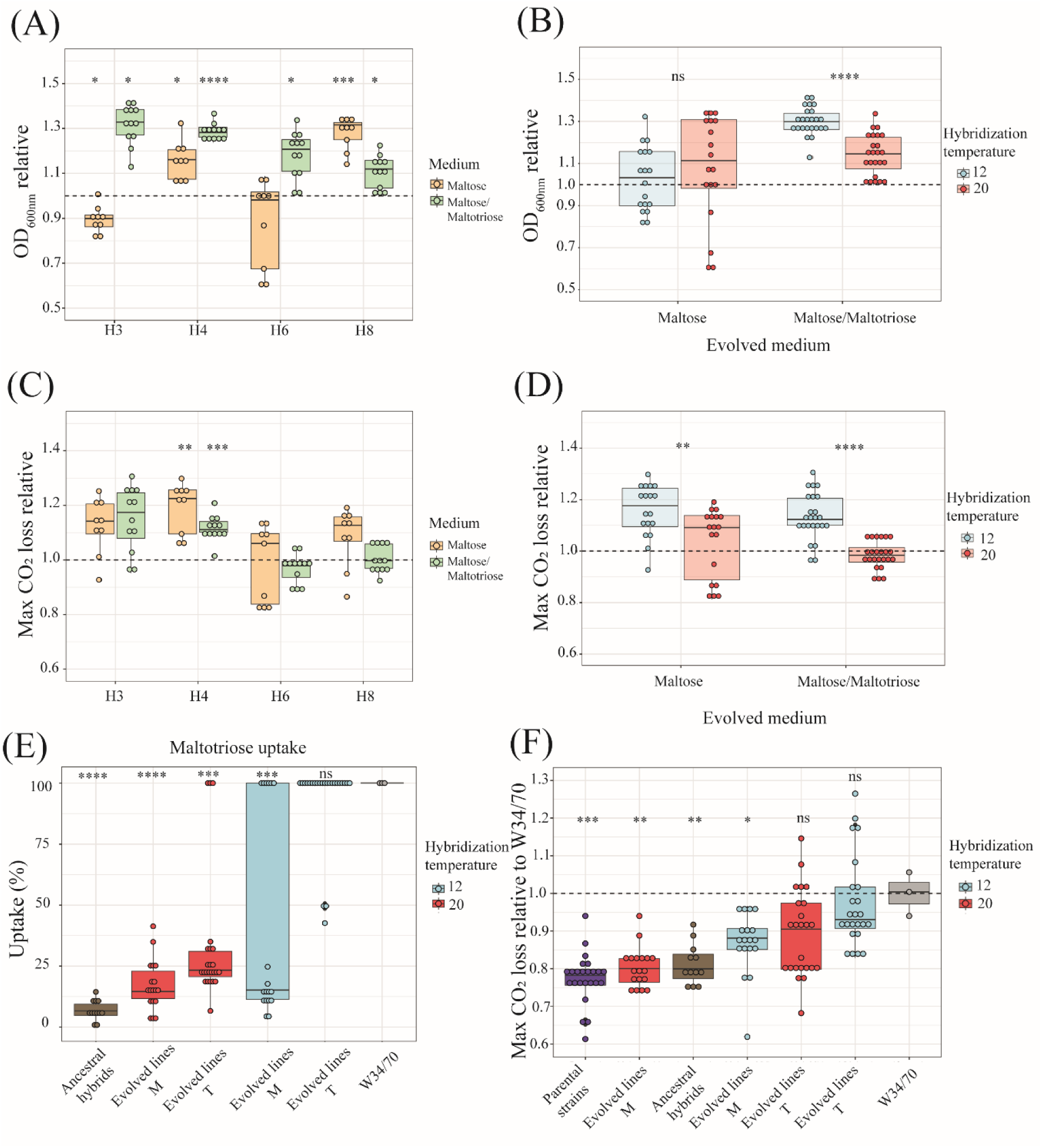
Fitness of evolved lines under microcultures and fermentation conditions. (A) Mean relative fitness (maximum OD_600nm_) of evolved lines after 250 generations under microculture conditions. Evolved lines were evaluated in the same medium where they were evolved (M or T medium). (B) Comparison of mean relative fitness (maximum OD600nm) shown in (A) between evolved lines from hybrids generated at 12 °C vs 20 °C. (C) Mean relative fitness (maximum CO2 loss) of evolved lines after 250 generation under fermentation conditions at 12 °C. (D) Comparison of mean relative fitness (maximum CO2 loss) shown in (C) between evolved lines from hybrids generated at 12 °C vs 20 °C. (E) Maltotriose uptake of evolved hybrid lines in maltose (M) and maltose/maltotriose (T), relative to the commercial Lager strain W34/70. Ancestral hybrids are shown in grey, cold-generated and warm-generated hybrid lines are shown in blue and red, respectively. (F) Fermentative capacity of evolved individuals relative to the commercial Lager strain W34/70 grouped according to the environmental condition used during experimental evolution and hybridization temperature to generate the ancestral hybrid. Plotted values correspond to the mean of three independent biological replicates of each evolved line or strain. Asterisk represents different levels of significance (Students t-test, * p ≤ 0.05, ** p ≤ 0.01, *** p ≤ 0.001, **** p ≤ 0.0001, ns not significant).

Next, we assessed the fermentative capacity of the evolved lines under conditions resembling beer wort fermentation (12 °Brix and 12 °C) (**Figure 2C, Figure S6,** and **Table S6A-S6B**). We did not observe a significant increase in CO_2_ production levels in the evolved lines of the H6-A and H8-A hybrids in either M or T media (**Figure 2C, Figure S6** and **Table S6B**, p-value < 0.05, one-way ANOVA). However, we found a significant greater CO_2_ production in the evolved lines of H4-A, evident in both evolution media, indicative of higher fermentation activity. The evolved lines of H3-A under T media also demonstrated a slightly higher CO_2_ production (**Figure 2C** and **Table S6B**, p-value < 0.05, one-way ANOVA, for H4 evolved lines and p-values of 0.0708 and 0.05149 for H3 evolved lines in M and T, respectively). Thus, both evolved hybrids lines generated at cold-temperature, carrying *S. eubayanus* mitochondria, showed a greater increase in CO_2_ production than hybrids carrying the *S. cerevisiae* mitochondria (**Figure 2C**). Specifically, hybrids with *S. eubayanus* mitochondria increased their maximum CO_2_ loss by 10.6% when evolving in M medium (p-value = 0.003698, one-way ANOVA) and by 13% in T medium (p-value = 1.328e-08, one-way ANOVA) (**Figure 2D**). This was predominantly due to an elevated maltotriose uptake (**Figure 2E** and **Table S6C**). Notably, the fermentative capacity of these hybrids reached that of the commercial strain (**Table S6D**, p-value > 0.05, one-way ANOVA). These findings strongly suggest that lines derived from hybrids generated at colder temperatures carrying *S. eubayanus* mitochondria and evolved in a complex maltose/maltotriose medium (T), significantly enhanced their lager fermentative capacity due to an increase maltotriose uptake during beer wort fermentation.

### Isolation of evolved genotypes with improved fermentative capacity and maltotriose uptake

To isolate individual representatives from the evolved population lines, we obtained one single genotype from each of the four hybrid lines at 250 generations (28 genotypes in total), which were then subjected to phenotypic evaluation in beer wort. These individual genotypes showed similar fermentation profiles as the population-level analyses above (**Figure 2F)**. Evolved hybrid genotypes carrying *S. eubayanus* mitochondria and evolved in T medium (maltose/maltotriose, H3-E and H4-E), showed higher CO_2_ production compared to H6-E and H8-E (p-value < 0.05, ANOVA, **Figure S7**). The genotypes with the largest significant fitness increase were derived from line H3-3 and H3-4 evolved in T conditions (**Figure S7**), which exceeded the commercial strain. Interestingly, two genotypes deriving from H6-A (carrying the *S. cerevisiae* mitochondria) evolved in T medium also showed a CO_2_ loss similar to the commercial strain (p-value = 0.90372, one-way ANOVA).

To focus more in-depth on the evolved lines with the highest fermentative capacity and carrying the *S. eubayanus* mitochondria (H3-4 and H4-1 evolved in T medium, **Figure S7**), we isolated three colonies from each of these two lines to evaluate their fermentative capabilities. Notably, the CO_2_ loss kinetics among these genotypes were comparable (p-value > 0.05, one-way ANOVA), with genotype #1 from line H3-4 exhibiting the highest CO_2_ loss (**Figure 3A**). All these genotypes’ fermentation profiles closely resembled that of the W34/70 commercial Lager strain, underscoring the significantly high fermentative capacity of these novel hybrids (p-value > 0.05, one-way ANOVA, **Figure 3A**). All genotypes consumed the maltotriose in the T medium completely (**Table S7A**), and ethanol production ranged from 3.50% to 3.78% v/v (**Figure 3B**), which is similar to the commercial strain (p-value > 0.05, one-way ANOVA). One genotype (H4-1-C3) showed a remarkable 7.1% increase in ethanol production compared to the commercial strain (**Figure 3B**, p-value = 0.001, one-way ANOVA).

**Figure 3.**
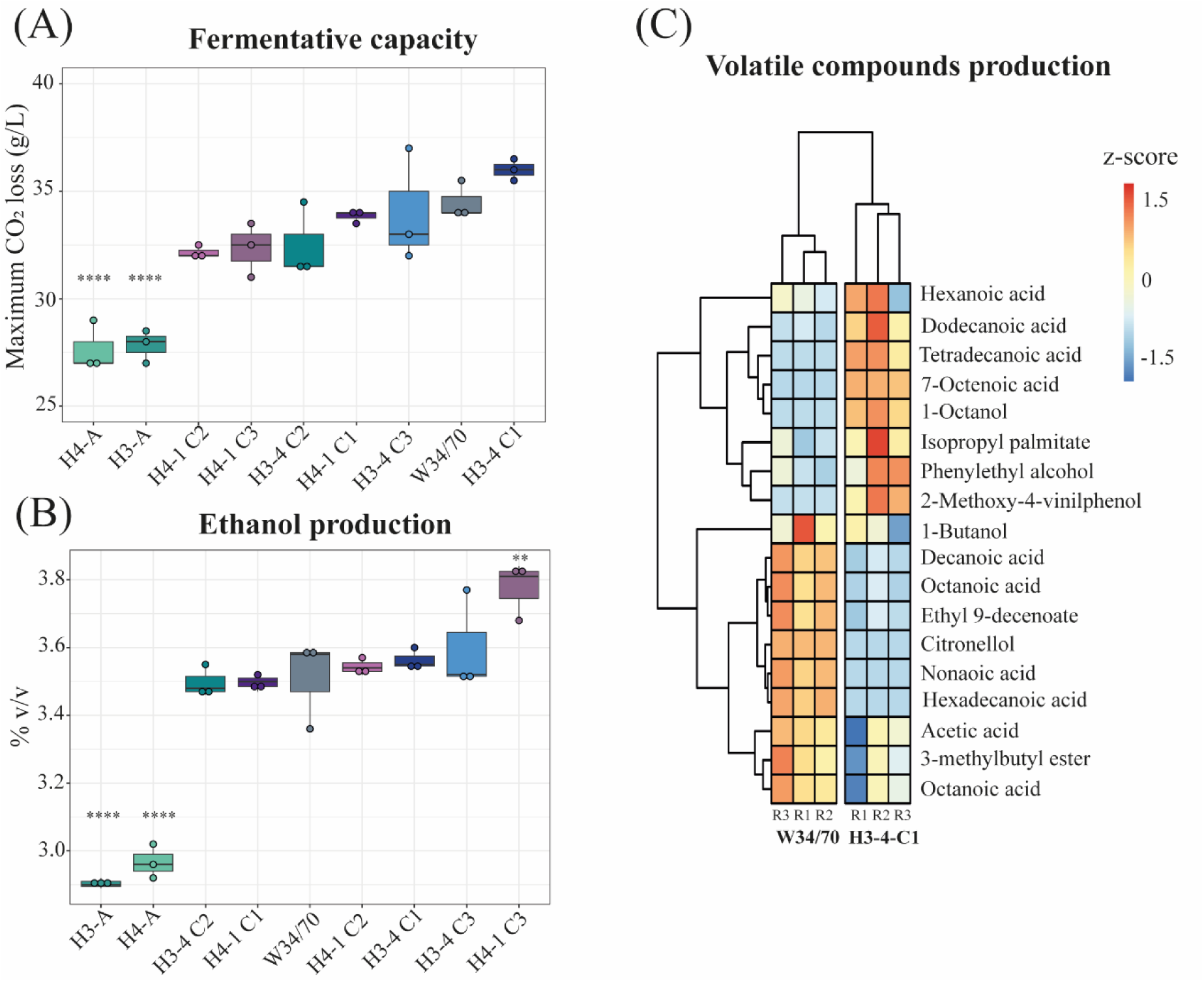
Fermentation performance of evolved hybrid individuals. (A) Maximum CO_2_ loss (g/L) for three different isolated genotypes (C1-C3) from evolved lines H3-4 and H4-1, ancestral hybrids (H3-A and H4-A) and commercial lager strain (W34/70). (B) Ethanol production (% v/v) for strains evaluated in (A). (C) Hierarchically clustered heatmap of volatile compounds production for strains evaluated in (A). Phenotypic values are calculated as normalized z-scores. For (A) and (B), plotted values correspond to the mean of three independent replicates. The (*) represents different levels of significance between hybrids and commercial lager strain (Student t-test, ** p < 0.01, **** p < 0.0001).

To compare the aroma profile of the H3-4-C1evolved hybrid to the lager strain, we identified volatile compounds (VCs) by HS-SPME-GC-MS in the fermented wort. This assay allowed us to identify 15 and 14 compounds in the evolved and commercial lager strains, respectively. We observed significant differences for 11 different compounds (**Figure 3C**, p-value < 0.05, one-way ANOVA, **Table S7B**), including ethyl esters and higher alcohols. For example, the evolved strain produced significantly more fatty acid ethyl ester, such as dodecanoic and tetradecanoic acid ethyl esters (p-value = 0.013 and 0.002, respectively, one-way ANOVA). The commercial lager strain on the other hand produced higher amounts of other ethyl esters, such as octanoic acid and nonanoic acid (**Figure 3D**, p-value = 0.001 and 0.0001, respectively, one-way ANOVA). The H3-4-C1evolved hybrid produced detectable levels of 4-vinyl guaiacol, which was completely absent in the lager strain. These results demonstrate that the aroma profiles of the evolved hybrids differ from the commercial lager strain.

### Evolved hybrids have mutations in genes related to carbon metabolism

To identify mutations in evolved hybrids associated with their improved fermentative capacity, we sequenced the genomes of the two genotypes exhibiting the highest CO_2_ production levels, specifically H3-4-C1 and H4-1-C1 (from here on referred to as H3-E and H4-E; with ‘E’ for evolved hybrid) that were evolved in the maltose/maltotriose T medium (**Table S8A**). Genome sequencing revealed that these two backgrounds had equal contributions from both parental genomes, that they had euploid, diploid genomes with no detectable aneuploidies (**Table S8B**), and that they contained *S. eubayanus* mitochondria.

We then identified *de novo* single nucleotide polymorphisms (SNPs) in the evolved hybrid genomes that were absent in the ancestral hybrids. We found 54 and 16 SNPs in the H3-E and H4-E backgrounds, respectively (**Table S8C**). The evolved hybrids differed in the total number of SNPs per genome (**Figure 4A**). In H3-E, we found 23 and 31 SNPs in the *S. cerevisiae* and *S. eubayanus* parental genomes, respectively, while H4-E only had 10 and 6 SNPs in the corresponding parental genomes. A GO-term analysis identified that many mutations in H3-E impacted genes related to ‘maltose metabolic process’, while in H4-E mostly ‘fungal-type cell wall organization’ genes were hit (**Figure S8**). We detected 27 SNPs within coding genes, of which six were related to maltose metabolism in the H3-E hybrid (**Table S8C**). For example, we identified an anticipated stop-codon in the *IRA2* allele (encoding for a GTPase-activating protein, **Figure 4B**) in the *S. cerevisiae* sub-genome, and a missense mutation in *MAL32* (encoding for a maltase enzyme) and *SNF3* (encoding for a plasma membrane low glucose sensor) in the *S. eubayanus* sub-genome (**Table S8C**), which are all genes related to sugar consumption.

**Figure 4.**
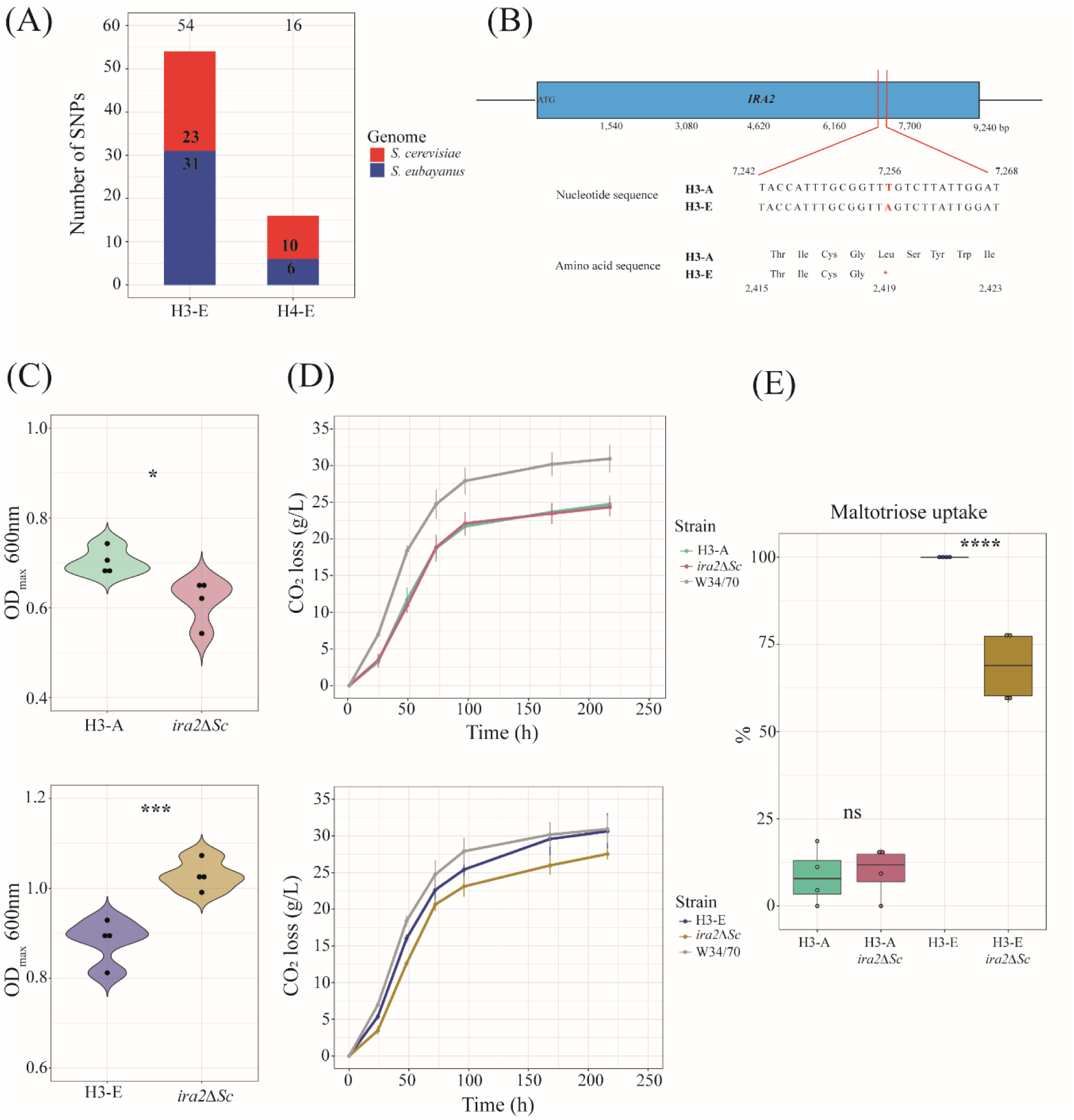
Genomic analysis of evolved hybrids. (A) Total number of *de novo* SNPs in the H3-E and H4-E hybrids. (B) SNP present in the *IRA2* gene in the *S. cerevisiae* sub-genome in the H3-E hybrid. (C) Maximum OD_600nm_ of *ira2Δ^Sc^* mutant strains under microculture conditions. Mutant and wild-type strains were evaluated in the T medium. (D) CO_2_ loss kinetics for *ira2Δ^Sc^* mutant and wild-type strains. (E) Maltotriose uptake (%) for strains evaluated in (D). For (C), (D) and (E), plotted values correspond to the mean of four independent replicates. The (*) represents different levels of significance between mutant and wild-type strains (Student t-test, * p < 0.05, *** p < 0.001, **** p < 0.0001).

To track the relative frequencies of the *IRA2, MAL32* and *SNF3* polymorphisms in the H3 evolution line, we sequenced whole population samples at increasing timepoints of experimental evolution (at 50, 100, 150, 200, and 250 generations; **Figure S9**). The *IRA2*-L2418* polymorphism arose before generation 50 and was completely fixed by 150 generations. Conversely, *MAL32* (K99N and L100Q) and *SNF3* (A493T) mutations only occurred late in evolution, between 200 and 250 generations, at low frequencies (10%).

To determine the phenotypic impact of the stop-codon detected in the *IRA2* gene, we performed a CRISPR assay targeting the *S. cerevisiae IRA2*, generating null mutants (*ira2^Sc^*) in the evolved and non-evolved hybrids. We evaluated growth under microculture conditions in the same evolutionary medium (T) and under beer wort fermentation (**Figure 4C**). This assay revealed that *ira2^Sc^* mutants in the H3-A hybrid background had a 12.5% lower OD_max_ under maltose/maltotriose conditions compared to H3-A (**Figure 4C**, p-value = 0.01213, one-way ANOVA), but still a similar fermentative capacity (**Figure 4D**, p-value = 0.79685, one-way ANOVA). In the H3-E hybrid, the null *ira2^Sc^* mutant showed a 16.6% higher OD_max_ under microculture conditions (**Figure 4C**, p-value = 0.00042, one-way ANOVA) and a significantly lower fermentative capacity under beer wort, with a 13.6% decrease in CO_2_ production (**Figure 4D**, p-value = 0.02315 one-way ANOVA, **Table S9**) and a 10.8% decrease in the maximum CO_2_ loss rate (**Table S9**, p-value = 0.02268 one-way ANOVA). This decrease in the fermentative capacity in the H3-E null mutant correlates with a lower maltotriose uptake (68.6%, **Figure 4E**). These results suggest that the stop-codon in *IRA2* in the evolved hybrids does not necessarily lead to a loss of protein function, but instead to a complex genetic interaction in the H3-E background promoting a trade-off between biomass and fermentative capacity, which is likely partly responsible for the phenotypic differences during the evolutionary process.

### Copy number variants of genes related to maltose metabolism are associated with improved fermentative capacity in evolved hybrids

Since *ira2* null mutants did not restore the full increase in fermentative capacity of the evolved hybrids, we examined genes exhibiting copy number variation (CNVs) in H3-E and H4-E hybrids (**Figure 5A**, **Table S8D**). Both H3-E and H4-E hybrids contained changes in copy number, particularly in the *MAL* gene family (**Figure 5A**, **Table S8D)**. For example, we identified 2 and 4 extra copies of the *MAL13* and *MAL11* genes in H4-E and H3-E, respectively.

**Figure 5.**
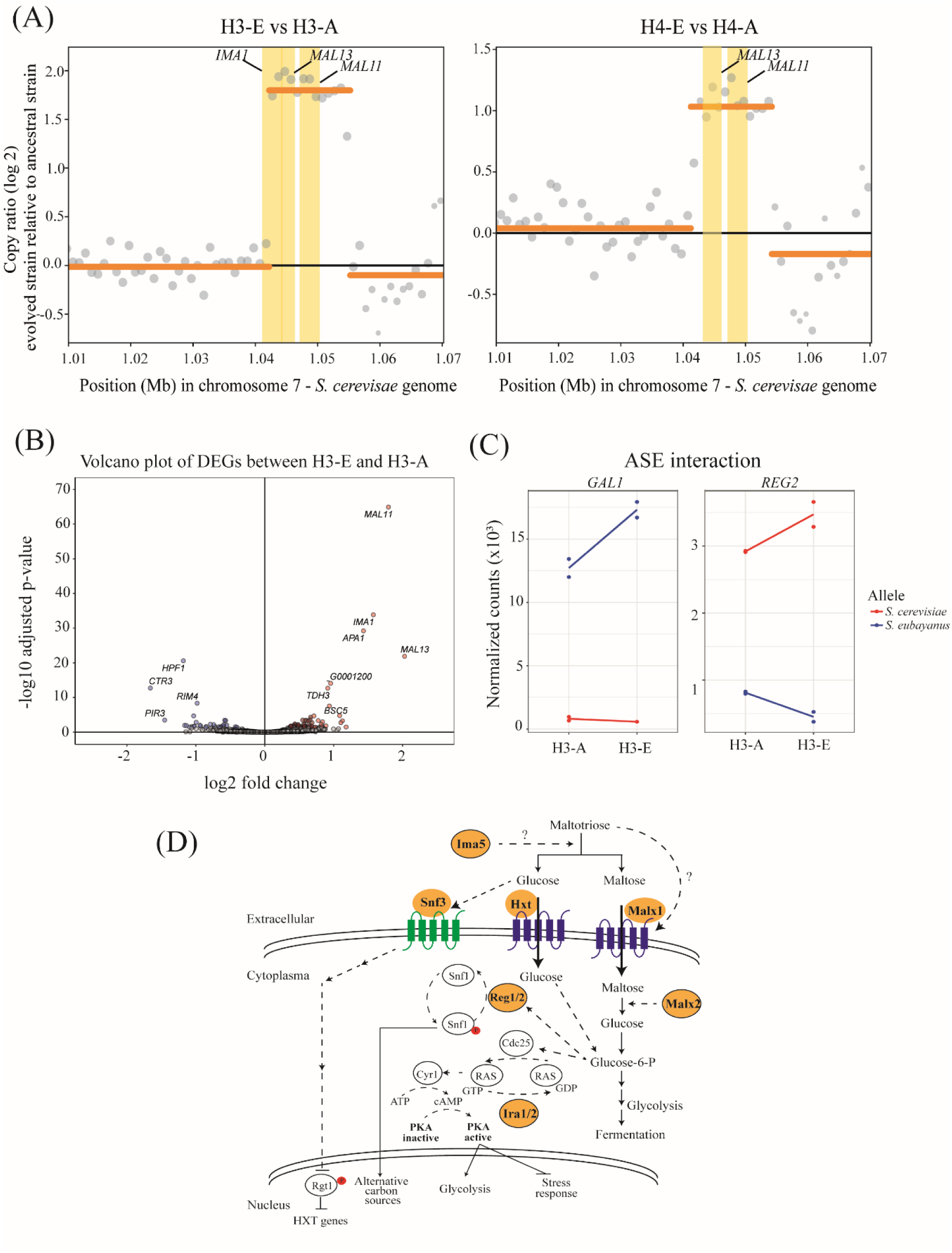
Copy number variation and differential gene expression analysis. (A) Copy number variations (CNVs) between H3-E and H4-E hybrids relative to their ancestral hybrids found in *S. cerevisiae* chromosome 7. Coding genes located within bins showing CNV calls higher than 1 copy (yellow rectangles) are shown. **(B)** Volcano plot showing differential expressed genes (DEGs) between H3-E and H3-A hybrids. The red and blue dots represent up-regulated and down-regulated genes in the H3-E hybrids, respectively. **(C)** Orthologous genes showing an interaction between allelic expression and experimental evolution. (D) Model depicting genes exhibiting mutations after the experimental evolution assay (highlighted in orange) and involved in pathways related to the detection, regulation, uptake, and catabolism of maltotriose. Phosphorylation is indicated in red. Green depicts a glucose sensing protein, while proteins in blue highlight transporters involved in sugar consumption.

To determine the impact of these mutations and the CNVs in the transcriptome of the H3-E hybrid, we estimated transcript abundance under beer fermentative conditions in the evolved and non-evolved hybrid. We identified 40 Differentially Expressed Genes (DEGs, FDR < 5%, **Table S8E**), where 21 and 19 genes were up- and down regulated in the evolved hybrid relative to its hybrid ancestor, respectively. Interestingly, we found that *S. cerevisiae* alleles for *IMA1*, *MAL11*, and *MAL13* were up-regulated in H3-E, which correlates with the increased gene copy number (**Figure 5B**). A GO term analysis showed that genes involved in maltose metabolic processes were up-regulated and genes in cell wall organization were down-regulated in the evolved hybrid, which correlates with the genetic changes we identified in coding regions (**Table S8F**).

To measure the impact of *cis*-variants on allelic expression within each parental subgenome, we estimated allele specific expression (ASE) in the evolved and non-evolved hybrids (**Figure 5C, Table S8G**). Seven genes showed ASE differences between the evolved and ancestral hybrid, likely originating from mutations in regulatory regions acquired during experimental evolution (**Figure 5C**, **Table S8G**). Of these, one and six ASE differences in the H3-E hybrid represented up-regulated alleles in the *S. cerevisiae* and *S. eubayanus* subgenomes, respectively (**Table S8G**). Interestingly, we detected the up-regulation of the *REG2* allele related to sugar consumption with a 2.1 higher fold change in the *S. cerevisiae* subgenome, which is involved in regulation of glucose-repressible genes (**Figure 5C**), correlating with the higher maltose and maltotriose consumption levels in the evolved hybrid.

## DISCUSSION

The hybrid yeast strains traditionally used for Lager beer production (*S. pastorianus*) are highly limited in genetic diversity. Currently, only two types of strains are used worldwide (Bonatto, 2021; Gallone et al., 2019; Gorter De Vries, Pronk, et al., 2019; Langdon et al., 2019), stemming from a single hybridization event that gave rise to the current Lager strains. This strongly constrains the diversity of available flavour and aroma profiles. The genetically depleted landscape of Lager strains also prevents a comprehensive understanding of the genetic changes crucial for the domestication process (Gallone et al., 2019; Langdon et al., 2019). The recent discovery of genetically and phenotypically distinct *S. eubayanus* lineages, including isolates from Patagonia (Eizaguirre et al., 2018; Langdon et al., 2020; Libkind et al., 2011; Nespolo et al., 2020), opened new avenues for understanding Lager yeast domestication, and to develop new strains to increase the diversity of fermentation profiles (Cubillos et al., 2019; Gibson et al., 2017). Previous studies using hybridization and experimental evolution demonstrated that Lager yeast hybrids could be improved through selection under fermentation conditions (Krogerus, Preiss, et al., 2018) without polyploidization (Krogerus et al., 2015). While these studies have expanded the diversity of Lager yeast phenotypes, they are primarily based on a single *S. eubayanus* genetic background, CBS 12357T (belonging to PB-1), which is not representative of the overall species’ genetic and phenotypic diversity (Burini et al., 2021; Molinet, Eizaguirre, et al., 2022; Nespolo et al., 2020; Urbina et al., 2020). *S. eubayanus* lineages vary widely in fermentation capacity and aroma profiles during beer fermentation, suggesting that the natural diversity of *S. eubayanus* is also well-suited for making innovative Lager hybrid strains (Burini et al., 2021; Mardones et al., 2021; Urbina et al., 2020). Here, we expanded the strain repertoire available for Lager brewing by including all three different *S. eubayanus* lineages found in Patagonia. Leveraging the genetic diversity of *S. eubayanus*, we created novel *S. cerevisiae* x *S. eubayanus* hybrids and enhanced their fermentation capacity through experimental evolution. We show that desirable phenotypic outcomes such as high ethanol production and new aroma profiles are the result of an intricate interplay of pre-existing genetic diversity, and selection on species-specific mitochondria, *de novo* mutations in sugar consumption genes, together with CNV of the *MAL* genes, important to improve maltose consumption during fermentation.

Hybridization offers a mechanism to combine beneficial traits from different species, which can enable adaptation to new environmental conditions (Gabaldón, 2020; R. Stelkens & Bendixsen, 2022) and improve the yield of plant cultivars and animal breeds (Adavoudi & Pilot, 2022; Rieseberg et al., 2003; Seehausen, 2004). However, hybridization - without subsequent selection of desirable traits for multiple generations - may not be sufficient to generate new phenotypes. None of the initial F1 hybrids in our experiment showed best parent heterosis, demonstrating that hybridization at different temperatures alone, was not sufficient to generate hybrids with a greater fitness than their parents under fermentative conditions. We therefore turned to experimental evolution as an alternative approach to improve the hybrids’ fermentative profiles. Experimental evolution across multiple generations, paired with time-series whole genome sequencing, is a powerful tool for studying microbial responses to a selective environment and to understand the fitness effects of *de novo* mutations (Barrick & Lenski, 2013; Burke, 2023; Cooper, 2018; Maddamsetti et al., 2015). We found that, after 250 generations in a high sugar and ethanol environment, hybrids evolved faster fermentation performance and higher ethanol production compared to both parents and ancestral unevolved hybrids. Interestingly, the hybrids’ evolutionary potential relied on the parental mitochondria. Hybrids with *S. eubayanus* mitochondria demonstrated higher fitness post-experimental evolution than those with *S. cerevisiae* mitochondria. Consistent with our results, all Lager commercial hybrids have *S. eubayanus* mitochondria (Gallone et al., 2019; Gorter De Vries, Pronk, et al., 2019; Langdon et al., 2019). It has been demonstrated that in synthetic hybrids, *S. eubayanus* mitochondria confers vigorous growth at colder temperatures compared to the *S. cerevisiae* mitotype, potentially conferring a competitive advantage in the cooler brewing conditions typical of Lagers (Baker et al., 2019). However, we performed experimental evolution at warmer temperatures (25°C). It is thus plausible that species-specific mitochondrial effects play an additional role, specifically concerning sugar utilization and glucose repression (Ulery et al., 1994), when adapting to Lager brewing conditions. These mitochondrial effects likely involve complex genetic interactions with the nuclear genome and might be exacerbated in the presence of the *S. eubayanus* mitochondria.

Our genome-wide screens for mutations to elucidate the genetic basis of hybrid fitness improvement identified several *de novo* SNPs and CNVs in the genomes of the evolved hybrids. These genetic changes were identified in genes with known effects on maltose metabolism and cell wall organization (**Figure 5D**). Particularly interesting are mutations in *IRA2* in the *S. cerevisiae* subgenome and in *SNF3* in the *S. eubayanus* sub-genome, which are both related to carbon metabolism. Evolved hybrids carried a premature stop codon in the *IRA2* gene, which was absent in both the *S. cerevisiae* and *S. eubayanus* parental ancestors, and in the unevolved hybrids at the beginning of experimental evolution. *SNF3* encodes for a low-glucose sensor and regulates the expression of hexose transporters (Santangelo, 2006), where *snf3* yeast mutants have low fitness in environments with low glucose concentrations (Vagnoli & Bisson, 1998). *IRA2* is a known suppressor of *snf3* mutants (Ramakrishnan et al., 2007), with *IRA2* required for reducing cAMP levels under nutrient limited conditions, where cAMP directly regulates the activity of several key enzymes of glycolysis (François & Parrou, 2001; Ramakrishnan et al., 2007). A mutation in *IRA2* would increase the carbon flux through glycolysis, which is in agreement with our finding that evolved hybrids showed higher sugar consumption. Furthermore, the regulation of the yeast mitochondrial function in response to nutritional changes can be modulated by cAMP/PKA signalling (Leadsham & Gourlay, 2010), which might be exacerbated in strains carrying *S. eubayanus* mitochondria. We further consolidated this mechanism by CNV and transcriptome analyses, which detected several up-regulated genes related to maltose consumption in the evolved hybrid during fermentation. Furthermore, the newly generated hybrids exhibited a distinct volatile compound profile compared to the W34/70 Lager strain. This highlights the potential of wild Patagonian yeast to introduce diversity into the current repertoire of available Lager yeasts. Previous studies in laboratory-made Lager hybrids revealed genetic changes that significantly impacted fermentation performance and changed the aroma profile of the resulting beer, compared to the commercial Lager strain (Gibson et al., 2020; Krogerus, Preiss, et al., 2018).

In summary, our study expands the genetic diversity of Lager hybrids and shows that new *S. cerevisiae* x *S. eubayanus* hybrids can be generated from wild yeast strains isolated from Patagonia. We found that hybridization at low temperatures, selecting for the retention of *S. eubayanus* mitochondria, followed by experimental evolution under fermentative conditions, and selection on desirable traits (ethanol production and aroma profiles), can generate hybrid strains with enhanced fermentation capacities. We delineate how genetic changes within distinct subgenomes of the hybrids contribute to improved fermentation efficacy, specifically in the context of cold Lager brewing conditions. This opens up new opportunities for the brewing industry to alleviate current constraints in Lager beer production, and to expand the range of currently available Lager beer styles.

## ACKNOWLEDGMENTS

We thank Antonio Molina, José Ruiz, Kamila Urbina, Mirjam Amcoff, Elin Gülich and S. Lorena Ament-Velásquez for their technical help. We also acknowledge Fundación Ciencia & Vida for providing infrastructure, laboratory space and equipment for experiments. This research was partially supported by the supercomputing infrastructure of the National Laboratory for High Performance Computing Chile (NLHPC, ECM-02) and by SNIC through Uppsala Multidisciplinary Center for Advanced Computational Science (UPPMAX) under Project naiss2023-22-62.

## AUTHOR CONTRIBUTIONS

Conceptualization: J.M., F.A.C.; Investigation: J.M., C.A.V., F.A.C.; Methodology: J.M., J.P.N., F.I.S., F.A.C.; Software: J.M., C.A.V., P.V.; Formal Analysis: J.M., J.P.N., R.S., F.A.C.; Resources: J.M., P.V., R.S., R.F.N., F.A.C.; Visualization: J.M., J.P.N., C.A.V., P.V.; Original Draft Preparation: J.M., C.A.V., R.S., F.A.C. All authors have read and agreed to the published version of the manuscript.

## Supplementary information

**Figure S1. Generation of interspecific *S. cerevisiae* x *S. eubayanus* hybrids.** Experimental procedure designed to generate and identify interspecific hybrids at two different temperatures (12 and 20°C).

**Figure S2. Phenotypic characterization of *S. cerevisiae* parental strains.** (A) Fermentation performance of 15 *S. cerevisiae* strains. (B) Maximum OD reached of growth curves in maltotriose 2% under microculture conditions (C) Maltotriose uptake after growth in maltotriose 2% under microculture conditions. Plotted values correspond to three biological replicates. The (*) represents different levels of significance between the phenotype of haploid strains and their respective parental strain (t-test; *p ≤ 0.05, **p≤ 0.01, ***p≤ 0.001, ****p≤ 0.0001 and ns: non-significant).

**Figure S3. Fermentative capacity at 12 °C of each hybrid.** Each plot represents a different cross. The (*) represents different levels of significance between the phenotype of hybrids and their respective parental strain (t-test; *p ≤ 0.05, **p≤ 0.01, ***p≤ 0.001, ****p≤ 0.0001).

**Figure S4. Fitness comparison of evolved lines after 250 generations.** (A) Mean relative fitness (growth rate) of evolved lines after 250 generation under microculture conditions. Evolved lines were evaluated in the same medium where they were evolved (M o T medium). (B) Mean relative fitness (growth rate) comparison between evolved lines from hybrids generated at 12 and 20 °C. Plotted values correspond to the mean of three independent replicates of each evolved lines. The (*) represents different levels of significance between evolved lines and unevolved hybrid in (A) and from hybrids generated at 12 °C vs 20 °C in (B) (Students t-test, * p < 0.05, ** p < 0.01, ns not significant).

**Figure S5. Fitness dynamics of evolved lines in maltose and maltose with maltotriose.** (A) Mean relative fitness (growth rate and OD) of replicate population in 2% maltose. (B) Mean relative fitness (growth rate and OD) of replicate population in 1% maltose and 1% maltotriose. Plotted values correspond to the mean of three independent replicates of each evolved line.

**Figure S6. Fitness dynamics of evolved lines in maltose and maltose with maltotriose under fermentation condition.** (A) Mean relative fitness (maximum CO_2_ loss) of replicate population in 2% maltose. (B) Mean relative fitness (maximum CO_2_ loss) of replicate population in 1% maltose and 1% maltotriose. Plotted values correspond to the mean of three independent replicates of each evolved line.

**Figure S7. Fermentative capacity of evolved individuals.** Fermentative capacity of evolved individuals relative to the commercial lager strain W34/70. Plotted values correspond to the mean of three independent replicates of each individual. The (*) represents different levels of significance between strains and commercial lager strain (Students t-test, * p < 0.05, ** p < 0.01, *** p < 0.001).

**Figure S8. GO term enrichment for genes with *de novo* mutations.** (A) Enriched GO terms identified in genes with *de novo* mutations in H3-E. (B) Enriched GO terms identified in genes with *de novo* mutations in H4-E.

**Figure S9. Dynamics of molecular evolution.** Allele frequencies over time in H3-A line evolved in T medium. In different colours are highlighted SNPs in the genes *IRA2*, *MAL32* and *SNF3*.

## Table Legends

**Table S1.** (A) Strains used in this study. (B) Primers used in this study.

**Table S2.** (A) Phenotypic characterization of the *S. cerevisiae* strains under fermentation conditions (maximum CO_2_ loss). (B) Statistical analysis of fermentative capacity of *S. cerevisiae* strains. (C) Sporulation efficiency and spore viability for *S. cerevisiae* and *S. eubayanus* strains.

**Table S3.** (A) Phenotypic characterization of the 31 interspecific hybrids and parental strains under microculture conditions. (B) Statistical analysis of phenotypes under microculture conditions. (C) Best-parent heterosis in the 31 interspecific hybrids evaluated under microculture conditions. (D) Mid-parent heterosis in the 31 interspecific hybrids evaluated under microculture conditions.

**Tables S4.** (A) Fermentation capacity (maximum CO_2_ loss) of hybrids in 12 °Brix wort at 12 °C. (B) Statistical analysis of fermentative capacity of hybrids at 12 °C. (C) Best-parent heterosis for fermentative capacity. (D) Mid-parent heterosis for fermentative capacity. (E) Sugar consumption and ethanol production of four interspecific hybrids and parental strains. (D) Statistical analysis of maltotriose uptake and ethanol production.

**Table S5.** (A) Mean relative fitness (growth rate and OD) and statistical analysis of each of the evolved lines in maltose and maltose/maltotriose relative to unevolved hybrid. (B) Mean relative fitness (growth rate and OD) and statistical analysis of evolved hybrids in maltose and maltose/maltotriose relative to unevolved hybrid. (C) SNPs identified in the *COX3* gen. (D) Identity matrix derived from *COX3* gen multiple alignment.

**Table S6**. (A) Mean relative fitness and statistical analysis for maximum CO_2_ loss of each of the evolved lines in maltose and maltose/maltotriose relative to unevolved hybrid. (B) Mean relative fitness and statistical analysis for maximum CO_2_ loss of evolved hybrids in maltose and maltose/maltotriose relative to unevolved hybrid. (C) Maltotriose uptake and statistical analysis of evolved lines in maltose and maltose/maltotriose relative to commercial Lager strain W34/70. (D) Mean relative fitness and statistical analysis for maximum CO_2_ loss of evolved lines in maltose and maltose/maltotriose relative to commercial Lager strain W34/70.

**Table S7.** (A) Fermentative capacity, maltotriose uptake and ethanol production of evolved individuals of H3-A and H4-A hybrids. (B) Volatile compounds production of H3-4-C1 and W34/70 in beer wort.

**Table S8.** (A) Bioinformatics summary statistics. (B) Genomic contributions (%) from parental strains in the H3-E and H4-E hybrids. (C) SnpEff analysis of the novel polymorphisms in H3-E and H4-E. (D) CNV results comparing evolved hybrids with their ancestral hybrid. Only CNVs with 1 or more copies are listed. (E) RNA-seq analysis between H3-E and H3-A hybrids. (F) Enriched GO terms of hybrid genes showing differential expression between ancestral and evolved hybrids. (G) Genes exhibiting Allele-Specific Expression (ASE), with values approximating 1 indicating overexpression of S. cerevisiae alleles, and values close to 0 representing overexpression of S. eubayanus alleles..

**Table S9.** Fermentative capacity and maltotriose uptake of *ira2* mutants.

